# Bacteriophage activity against and characterisation of avian pathogenic *Escherichia coli* isolated from colibacillosis cases in Uganda

**DOI:** 10.1101/2020.09.01.277269

**Authors:** George Kazibwe, Phionah Katami, Ruth Alinaitwe, Stephen Alafi, Ann Nanteza, Jesca Lukanga Nakavuma

**Affiliations:** School of Biosecurity, Biotechnical and Laboratory Sciences, College of Veterinary Medicine, Animal Resources and Biosecurity, Makerere University. P.O. Box 7062, Kampala Uganda

**Keywords:** APEC, virulence genes, phylogenetic groups, antibiotic susceptibility, bacteriophage

## Abstract

A laboratory-based study aimed at establishing a stock of avian pathogenic *Escherichia coli* (APEC) lytic bacteriophages, for future development of cocktail products for controlling colibacillosis as well as minimizing use of antimicrobial drugs in the poultry production systems in Uganda. Specifically, the study determined the antibiotic susceptibility; phylogenetic categories, occurrence of selected virulence genes among *Escherichia coli* stock isolates from cases of chicken colibacillosis; and isolation of specific bacteriophages. Fifty six isolates were confirmed as *E. coli* by standard phenotypic tests. All the 56 (100%) isolates were resistant to at least one antibiotic while 50 (89.3%) isolates were resistant to at least three classes of antimicrobial drugs and were therefore designated as multi-drug resistant. Phylogenetically, APEC isolates mainly belonged to phylogroups A and D which represented 44.6% and 39.3%, respectively. Virulence genes, *ompT* and *iutA* were the most frequent with 33 (58.9%) and 32 (57.1%) isolates respectively; while *iroN* least occurred in 23 (41.1%) isolates. Of the 56 isolates, 69.6% harbored at least one virulence gene, while 50% had at least four virulence genes; hence confirmed as APEC. None of the isolates belonged to the selected serotypes O1, O2 and O78. Seven specific bacteriophages were isolated and their host range, varied from 1.8% to 17.9% (n=56 APEC isolates), while the combined lytic spectrum of all the phages was 25%. Phage stability was negatively affected by increasing temperatures with both UPEC04 and UPEC10 phages becoming undetectable at 70°C; however activity was detected between pH 2 and 12. The high occurrence of APEC isolates with resistance against the commonly used antibiotics supports the need for alternative strategies of bacterial infections control in poultry. The low host range exhibited by the phages calls for search for more candidates before more in-depth studies are done for phage characterization and application.

## 1. Introduction

Avian colibacillosis refers to any localized or systemic infection caused by Avian Pathogenic *Escherichia coli* (APEC) belonging to several serogroups; and remains one of the most prevalent bacterial diseases affecting the poultry industry worldwide [1]. In Uganda, colibacillosis is the most frequent bacterial infection among the chicken samples submitted to the Central Diagnostic Laboratory with a prevalence of 14% [2]. Majalija *et al* (2010) reported 87% *Escherichia coli*, isolated from broiler farms kept under the deep litter system, being resistant to at least one antimicrobial agent [3].

*Escherichia coli* strains possess various virulence factors for extra-intestinal survival [4– 8]. However, capacity to cause disease in a specific host species, depends on acquisition of appropriate virulence gene combination by a given *E. coli* strain [5]. The number of detected genes can be used as a reliable index of their virulence; and strains typed as APEC possess five to eight genes, while the non-APEC ones harbor less than four genes [5,8]. Virulence-associated genes, namely, *iutA, hlyF, iss, iroN*, and *ompT* were suggested as the minimum that can be used to identify an APEC strain with the highest pathogenicity [9].

Bacterial diseases of significance in animal production systems affect productivity, may be zoonotic and some have been associated with drug resistant pathogens [6,10]. The high occurrence of drug resistant organisms warrants search for alternative strategies, such as, use of bacteriophages, in management of bacterial infections, like colibacillosis. Bacteriophages are naturally occurring viruses in the environment that routinely control bacterial populations [11]. The specific action against bacteria, self-replicating and self-limiting nature; makes bacteriophages attractive alternatives to antibiotics to prevent and treat bacterial diseases. Phages have been applied in control of various bacterial agents; including the drug resistant strains. Bacteriophages are increasingly explored as non-antibiotic strategies for control of bacterial diseases in agricultural systems for food safety and security [12–14]. In some developed countries, phages have been approved and are commercially available for use, especially in management of microbial contamination of plant-based foods [15]. In poultry, phages have demonstrated effectiveness; and some e.g. ListShield™from Intralytix, Inc; have been approved by the United States Food and Drug Administration (FDA) for managing bacterial infections including *E. coli* (EcoShield™) [12,16,17].

The APEC strains circulating on poultry farms in Uganda have neither been characterized nor the virulence genes they harbor documented. Unlike most poultry diseases, there are no vaccines for controlling colibacillosis, hence its management depends on hygienic measures as well as use of antibacterial agents. Antibiotic use is associated with resistance development and undesirable drug residues in the poultry products; which calls for alternative antimicrobial strategies. The research aimed at establishing a stock of APEC lytic bacteriophages, for future development of cocktail products for controlling colibacillosis as well as minimizing use of antimicrobial drugs in the poultry production systems in Uganda. Specifically, the *E. coli* isolates associated with cases of poultry colibacillosis were characterized by drug susceptibility, phylogenetic group and the virulent genes harbored. The research also sought to establish presence of APEC serotypes O1, O2 and O78; and stock of lytic bacteriophages that specifically target APEC.

## 2. Materials and Methods

### 2.1 Bacterial isolates

Previously archived APEC isolates from post-mortem samples of colibacillosis suspect chicken collected between 2017 and 2018 from poultry farms around Kampala district were used for the study. The isolates had been stored at the microbiology laboratory of the College of Veterinary Medicine, Animal Resources and Biosecurity, Makerere University. Identity of 56 *Escherichia coli* isolates was confirmed by standard bacteriological and biochemical methods.

### 2.2 Antimicrobial susceptibility testing

Antimicrobial susceptibility testing was carried out by the Disk diffusion method [18], as recommended by the Clinical and Laboratory Standards Institute [19]. Twelve antibiotics, including Tetracycline 25mcg, Chloramphenicol 5mcg, Nalidixic acid 30mcg, Ampicillin 10mcg, Streptomycin 10mcg, Co-trimoxazole 25mcg, Ciprofloxacin 5mcg, Penicillin G 10mcg, Cefixime 30mcg, Amoxicillin 30mcg, Nitrofurantoin 300mcg, and Gentamicin 30mcg (Bioanalyse^®^) were tested on Mueller-Hinton agar. Growth-inhibition zones were recorded and interpreted as susceptible (S), intermediate (I), and resistant (R). An *E. coli* reference strain (ATCC 25922) was used for quality control of the test.

### 2.3 DNA extraction

Template DNA was extracted using the boiling method as described by Wang *et al* [20]. Briefly, bacteria DNA was prepared by suspending one colony of the isolate in 100μL of distilled water. The suspension was rapidly boiled in a water bath at 95°C for 10 minutes and then cooled to room temperature. The cool suspension was then centrifuged (Eppendorf centrifuge 5424R, Germany) for 3 minutes at 12000rpm to remove cell debris; and the supernatant stored at −20°C formed the stock from which aliquots of template DNA were obtained for use in PCR.

### 2.4 Detection of the virulence genes and determination of phylogroups and serogroups of APEC using PCR

Amplification of the selected *E. coli* virulence genes was carried out following a method described by Johnson *et al* [9]. The positive controls used in PCR assays were *E. coli* strains BEN2268, BEN2908 which were kindly provided by Dr. Catherine Schouler.

A triplex PCR was carried out following a method described by Clermont *et al* [21] to determine the phylogenetic groups of the APEC isolates; where four major phylogenetic groups (A, B1, B2 and D) were targeted. The *E. coli* K-12 (phylogroup A), STEC O111 (phylogroup B1), and O157:H7 (phylogroup D) were used as positive controls.

Serogroup identification was done using an allele-specific PCR assay with primers designed for the most common serotypes (O1, O2 and O78) as described by Wang *et al* [22]. *E. coli* strains BEN2268 and BEN2908 were used as positive controls with nuclease free water used as the negative control. The primers and the PCR conditions used are listed in the supplementary files (S1-S3 Appendices).

### 2.5 Isolation of bacteriophages

*Escherichia coli* specific phages were isolated through enrichment, from effluent and chicken droppings that were obtained from three selected chicken houses and slaughter places around Kampala district. The phage isolation process followed the procedure described by Oliveira *et al* [23] with slight modifications including use of Tryptic soy broth (TSB) (Condalab, Madrid, Spain) instead of Luria Bertani broth (LB). Briefly, 50g of the faecal samples were homogenized in 50 ml of Tryptic soy broth (TSB). The effluent (50 ml) and the homogenised samples were centrifuged at 10,000 ×g for 10 min. The supernatant was filtered through a 0.45μm membrane (ADVANTEC^®^, USA) and 10 ml of the filtrate was added to 10 ml of double strength TSB containing 40μL of 1M Calcium Chloride (CaCl_2_). Then 100μL of overnight *E. coli* ATCC 25922 broth culture was added for enrichment. The mixture was incubated at 30°C for up to 48 hours on a shaker (New Brunswick™ Innova^®^ 40, Germany) at 120 rev/min; after which it was centrifuged at 7000 rpm (Hermle Z32K, Germany) for 5 mins at 4°C. The supernatant was then filtered through 0.45μm syringe filters. Presence of phages was determined using the spot assay method.

### 2.6 Spot assay method

A spot assay was carried out as described by Mirzaei & Nilsson [24] with slight modifications. Briefly, the soft agar overlay was prepared by mixing 100 μL of an overnight *E. coli* broth culture with 5mL of TSB containing 0.7% agar maintained in the molten form in a water bath at 45°C. The agar overlay was poured on to base plates containing 20–30mL of Tryptic Soy Agar (TSA) (Condalab, Madrid, Spain) with 1.5% agar and then swirled to allow uniform spread. On solidifying, 10 μL of the phage filtrate was spotted on top of the soft agar and allowed to dry. The plates were examined for lysis or plaque formation after overnight incubation at 37°C. A clear zone indicated presence of phage.

### 2.7 Purification of bacteriophages

Phage purification was done using the agar overlay technique as described by Oliveira *et al* [23], with some modifications. The method employed base plates, containing 20–30mL of TSA with 1.5% agar and soft agar overlays composed of TSB with 0.7% agar. Ten-fold serial dilutions (10^0^ - 10^−9^) of the above filtered phages were prepared using the phage SM buffer (0.05M Tris, 0.1M NaCl, 0.008M MgSO_4_, 0.01% w/v gelatin, pH 7.5). Equal volumes (100 μL) of the diluted phage and of overnight host *E. coli* were mixed with 5mL of soft agar overlay, spread onto TSA plates and incubated overnight at 37°C. Phages were purified by successive single plaque isolation, from the higher dilutions plates where plaques were distinct. A single plaque was picked from the bacteria lawn, suspended into an overnight host *E. coli* culture, incubated overnight at 37°C and the lysate plated as described above. After repeating the cycle three more times, lysates from single plaques were centrifuged at 5000 g for 5 min. The phages were recovered from the supernatant by filtering through a 0.45 μm membrane. Purified phages were stored in SM buffer at 4°C for working stock, while for long term-storage, phage stocks were stored in 1 ml aliquots at −80°C in 7% Dimethyl Sulfoxide (DMSO).

### 2.8 Determination of phage titres by agar overlay method

Phage concentration (titre) was determined using a method described by Carey-Smith *et al* [25] with some modifications. Tryptic Soy broth instead of Luria Bertani broth was used as the culture medium. Ten-fold serial dilutions (10^0^ – 10^−8^) of the purified phages were prepared using SM buffer. Overlays (5ml) were inoculated with 100 μL of overnight host *E. coli* and poured on a base plate previously marked in a grid to allow identification of each phage dilution. Once the overlay was gelled and dried, 10 μL of each phage dilution was spotted. The plates were incubated at 37°C, and examined for plaques after 24 hours. Distinct plaques obtained from the lowest dilution were counted and used to calculate the phage titre. The titres were expressed as plaque forming units (PFU) per ml.

### 2.9 Bacteriophage host range determination

Bacteriophage activity was tested on 56 APEC isolates using the spot assay method as described above. Presence of clear zones indicated sensitivity of a given APEC isolate to the lytic activity of the phage. Out of the seven phages, two phages with the broadest host ranges were selected for pH and thermal stability testing.

### 2.10 pH and thermal stability test

pH stability and thermal stability tests were carried out for the two phages with the broadest host ranges as described by Jung *et al* and Yu *et al* [26,27]. Briefly, the phages (10^8^ PFU/ml) were incubated at different temperatures (20°C to 70°C) for 30 mins. This range of temperatures was selected because it encompasses both the room temperature and body temperature of chicken among other temperatures. Afterwards, the phage suspensions were immediately placed in an ice bath.

The pH stability of the phages was evaluated using SM buffer solution adjusted to the required pH using concentrated hydrochloric acid (HCl) or Sodium hydroxide (NaOH). The phages (10^8^ PFU/ml) were subjected to different ranges of pH from 2 to 12 for 30 mins at 25°C and at 40°C. The two temperatures were selected to represent the room temperature and the body temperature of chicken respectively; while pH was studied because it affects phage adsorption onto the bacteria and its subsequent propagation. Afterwards, the phage suspensions were immediately diluted with the SM buffer to limit further exposure. After both the heat and pH treatment, viable phages were quantified using the agar overlay method as described above. All assays were performed in duplicates.

### 2.11 Research approval

The study was endorsed by the Higher Degrees Research Committee of the College of Veterinary Medicine, Animal Resources and Biosecurity of Makerere University.

## 3. Results

### 3.1 Antimicrobial susceptibility testing

All the 56 (100%) isolates exhibited resistance to at least one antibiotic. Fig 1 presents the proportion of resistant isolates for each of the tested antibiotic. High frequency of resistance was encountered for the antibiotics: Penicillin G (100%), Sulphamethoxazole/Trimethoprim (87.5%), Tetracycline (83.9%), Ampicillin (80.4%), Amoxicillin (69.6%), Streptomycin (67.9%) and Nalidixic acid (60.7%). Average frequency of resistance was found in case of Chloramphenicol (35.7%). Low frequency of resistance was revealed in case of Gentamicin (10.7%) and Nitrofurantoin (8.9%); while all the 56 (100%) isolates were susceptible to Cefixime. Resistance to at least three antimicrobial drug classes; and hence multi-drug resistance (MDR), was encountered in 50 (89.3%) isolates.

**Fig 1.**
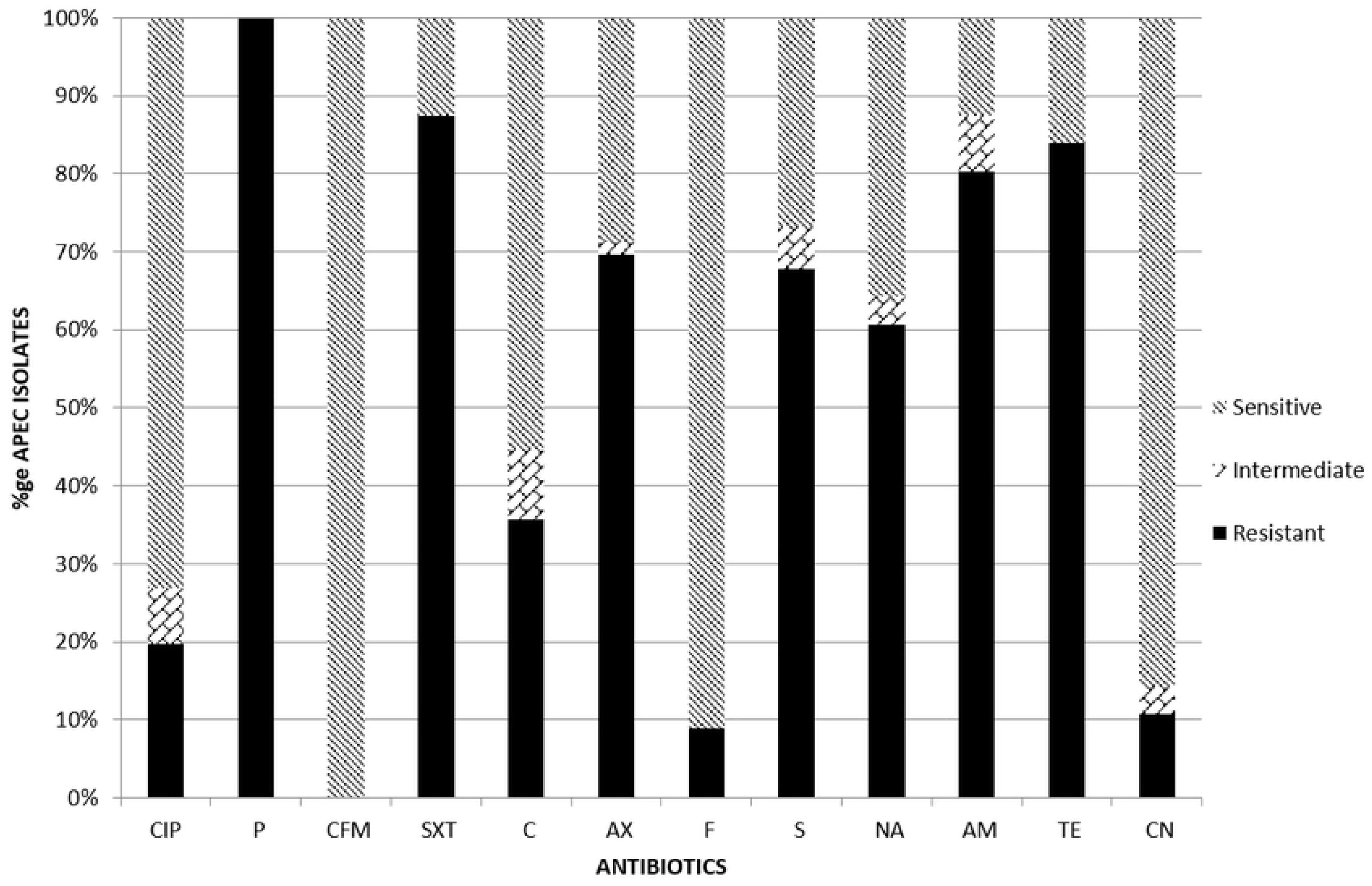
Antimicrobial susceptibility test results for Avian Pathogenic *E. coli*. The bars represent the percentages of the 56 APEC isolates that were resistant, intermediate or susceptible to the 12 antibiotics as determined by the Disk diffusion method.

CIP – Ciprofloxacin, P-Penicillin G, CFM – Cefixime, SXT - Sulphamethoxazole/Trimethoprim, C-Chloramphenicol, AX – Amoxillin, F – Nitrofurantoin; S – Streptomycin, NA - Nalidixic acid, AM - Ampicillin, TE-Tetracycline, CN – Gentamicin

### 3.2 Phylogenetic groups of the APEC isolates

The multiplex PCR amplification targeting the *ChuA, yjaA* and TspE4.C2 genes categorized the 56 APEC isolates into phylogenetic groups A, B1, B2 and D with 25 (44.6%), eight (14.3%), one (1.8%) and 22 (39.3%) isolates, respectively. Table 1 presents the genes and/or their combinations, the phylogenetic group and proportion of the *E. coli* isolates in each category.

**Table 1.**
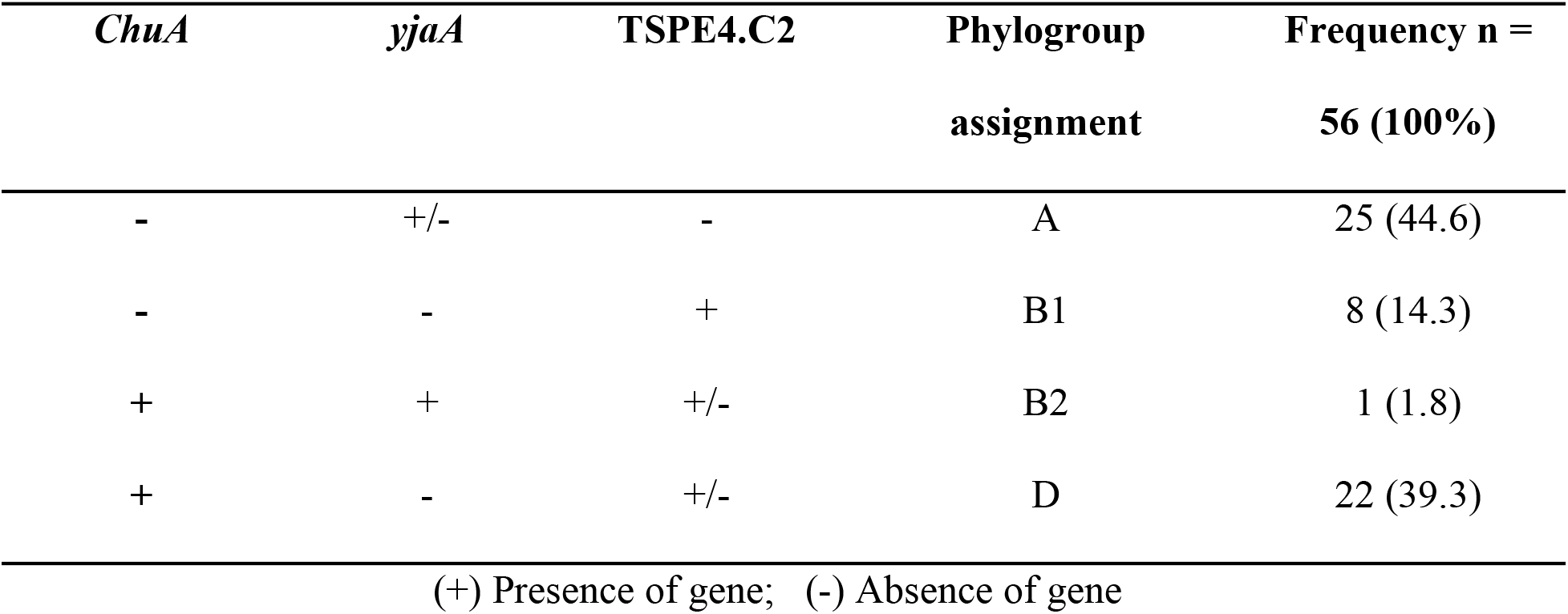
Phylogenetic groups of the APEC suspect isolates.

| *ChuA* | *yjaA* | TSPE4.C2 | Phylogroup<br>assignment | Frequency n =<br>56 (100%) |
| --- | --- | --- | --- | --- |
| - | +/- | - | A | 25 (44.6) |
| - | - | + | B1 | 8 (14.3) |
| + | + | +/- | B2 | 1 (1.8) |
| + | - | +/- | D | 22 (39.3) |

(+) Presence of gene; (-) Absence of gene

### 3.3 Frequency of APEC virulence genes

Table 2 presents the frequency of the APEC isolates harboring the selected virulence genes, that is, *iroN, ompT, hlyF, iss,* and *iutA.* Out of the 56 isolates, 39 (69.6%) had at least one virulence gene. The virulence genes *ompT* and *iutA* had the highest prevalence at 33 (58.9%) and 32 (57.1%) respectively with *iroN* having the lowest prevalence at 23 (41.1%). Of the 56 isolates, only 28 (50%) harboured four or more virulence genes and thus confirmed as APEC.

**Table 2.** Frequency of the selected virulence genes among the APEC suspect isolates.

| Gene | Description | Frequency<br>n = 56 (100%) |
| --- | --- | --- |
| *iutA* | Aerobactin siderophore receptor gene | 32 (57.1) |
| *iss* | Episomal increased serum survival gene | 31 (55.4) |
| *hlyF* | Putative avian hemolysin | 31 (55.4) |
| *ompT* | Episomal outer membrane protease gene | 33 (58.9) |
| *iroN* | Salmocheilin siderophore receptor gene | 23 (41.1) |

With respect to presence of at least one virulence gene in relation to the phylogroup; 14 out of 25 in group A, 5 out of 8 in group B and 19 out of 22 in group D, had virulence genes (Fig 2).

**Fig 2.**
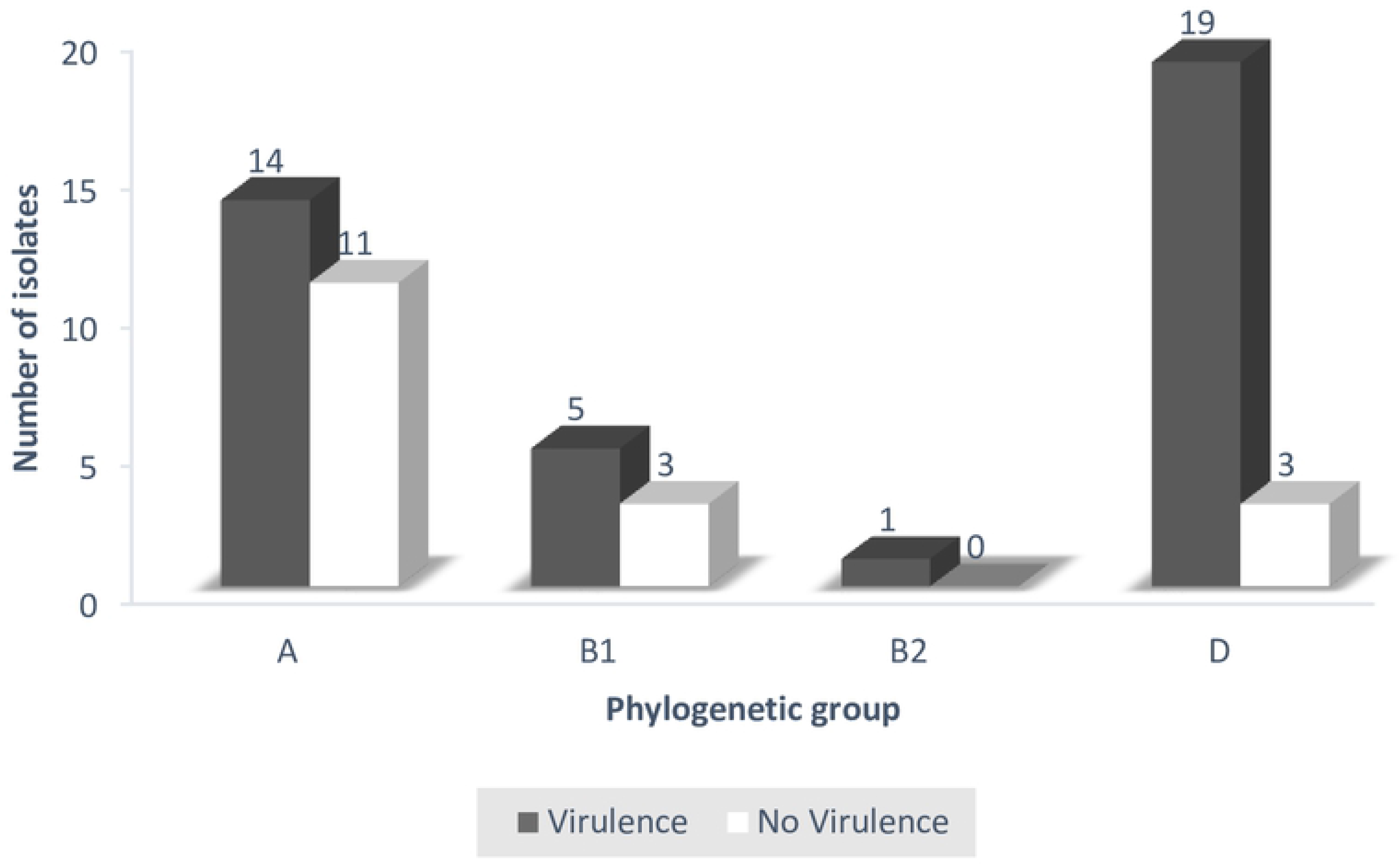
Virulence gene content of APEC isolates within each phylogenetic group. The dark bars indicate the proportion of APEC isolates within a phylogenetic group that had virulence genes while the white ones indicate those without virulence genes.

### 3.4 Serological genotyping

Of the 56 isolates, none generated amplicons of sizes expected for the O1, O2 and O78 serogroups (Fig 3).

**Fig 3.**
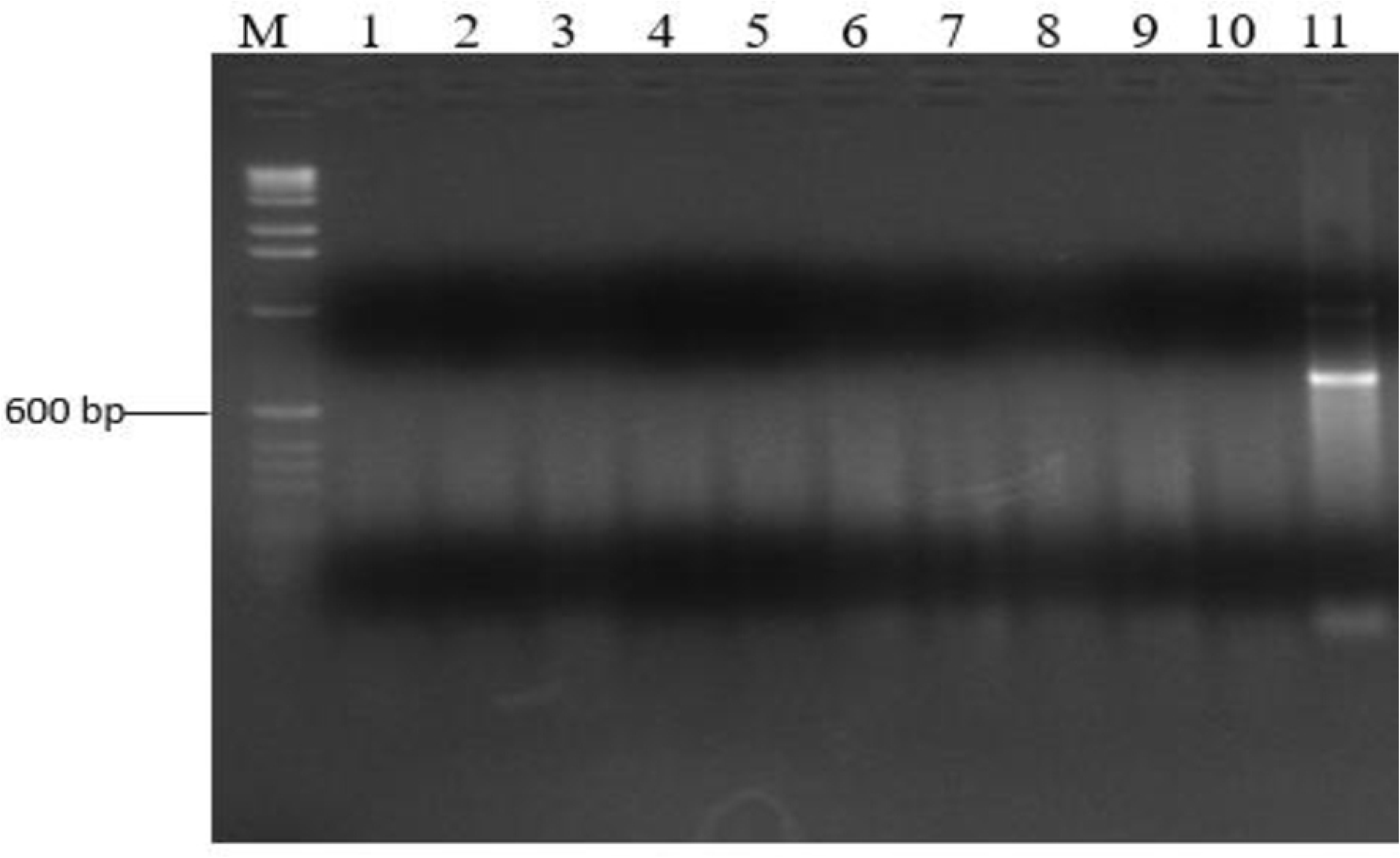
Agarose gel showing PCR amplicons from selected APEC isolates for Serogroup O78. Lane M: DNA marker (100bp DNA ladder, ThermoFisher Scientific); Lanes 1-10: APEC isolates; Lane 11: APEC strain BEN2268 (positive control).

### 3.5 Phage isolates and their host range

A total of 10 crude phage isolates were obtained but seven were successfully purified. The purified phages were code-named as UPEC01, UPEC03, UPEC04, UPEC06, UPEC08, UPEC09 and UPEC10. The phage host range, as exhibited by lytic activity against 56 APEC isolates varied from one (1.8%) to 10 (17.9%). Phage UPEC04 had the broadest host range, inhibiting 10 (17.9%) APEC isolates followed by UPEC06 and UPEC10 at 6 (10.7%) isolates each, then UPEC03 at 5 (8.9%) isolates, UPEC01 and UPEC08 at 4 (7.1%) isolates each; while UPEC09 had the narrowest host range of 1 (1.8%) isolate. Only 14 (25%) APEC isolates out of the 56 were sensitive to any one phage and the combined lytic spectrum of UPEC04 and UPEC10 phages includes all the total APEC isolates that were sensitive. Therefore UPEC04 and UPEC10 phages were selected for further analysis. Out of the 14 APEC isolates sensitive to the phages, 11 were multi drug resistant. The phage sensitivity pattern of the seven phages on the 14 APEC isolates is presented in S1 Table.

### 3.6 Thermal and pH stability of UPEC04 and UPEC10 phages

Phages UPEC04 and UPEC10 were selected for further investigation because the combined lytic activity of the two yielded the maximum host range of 14 out of the 56 tested APEC isolates. Therefore, the heat sensitivity of these two phages was determined for temperatures ranging from 20°C - 70°C (Fig 4). The phages were stable to heat with only slight reductions in titers up to 50°C, followed by a steep decline up to 70°C; beyond which they were undetectable. The highest titers were obtained between 20°C – 50°C making this the range of temperature at which the two phages are most stable.

**Fig 4.**
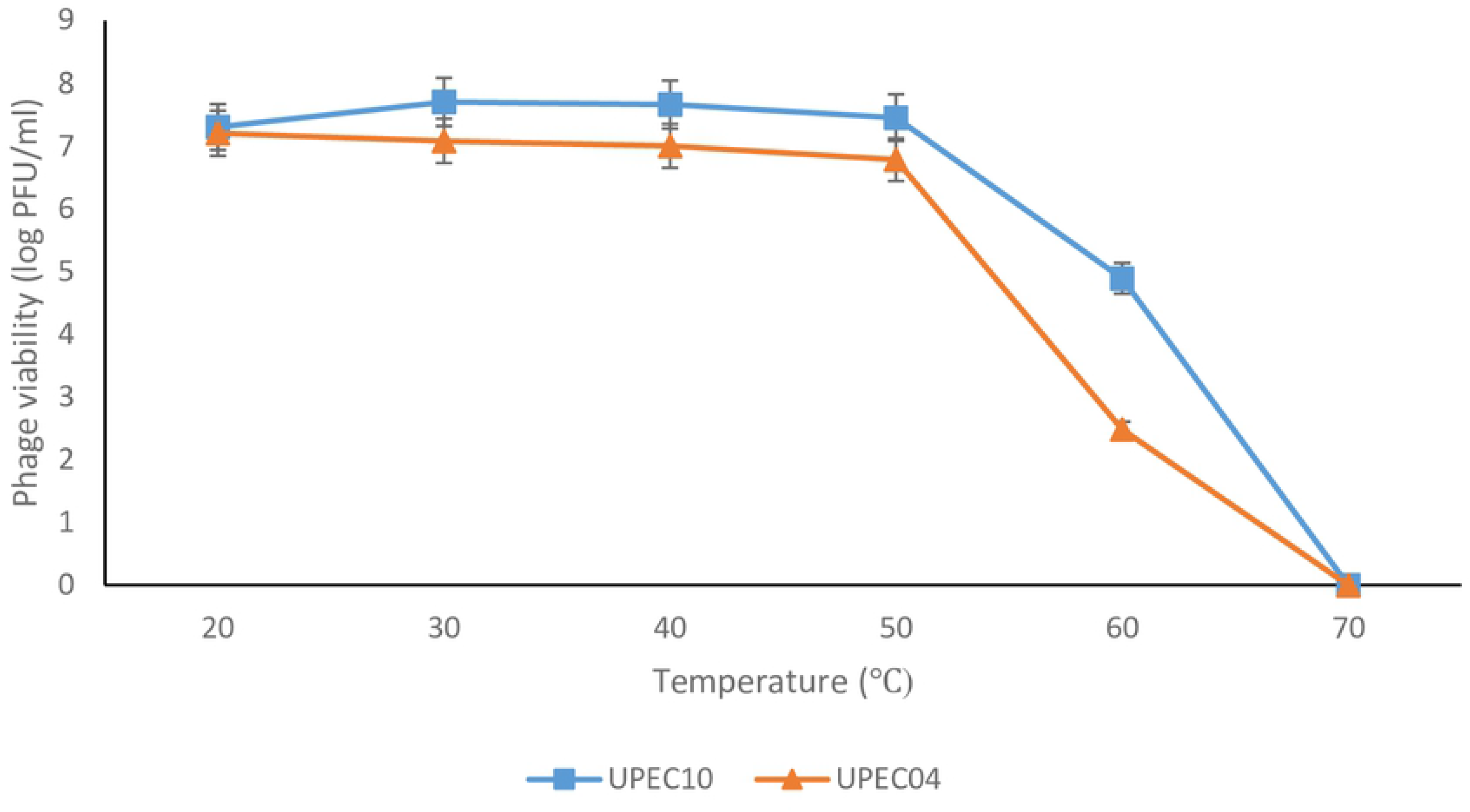
Effect of temperature on UPEC04 and UPEC10 phage viability. Phage viability was determined by obtaining the phage titers at the different temperatures using the agar overlay method. Values are an average for duplicate tests.

### 3.7 Effect of pH on phage titer

The stability of UPEC04 and UPEC10 to pH ranges from 2 to 12 at both 25°C and 40°C is presented in Fig 5. The phages retained viability across the different pH values with the lowest titers registered at the extremes of pH (2 and 12), while the highest titers were registered between pH 4 and 8. The changes in the titers followed a similar pattern at the two temperatures, though the titers were consistently higher at 25°C compared to 40°C.

**Fig 5.**
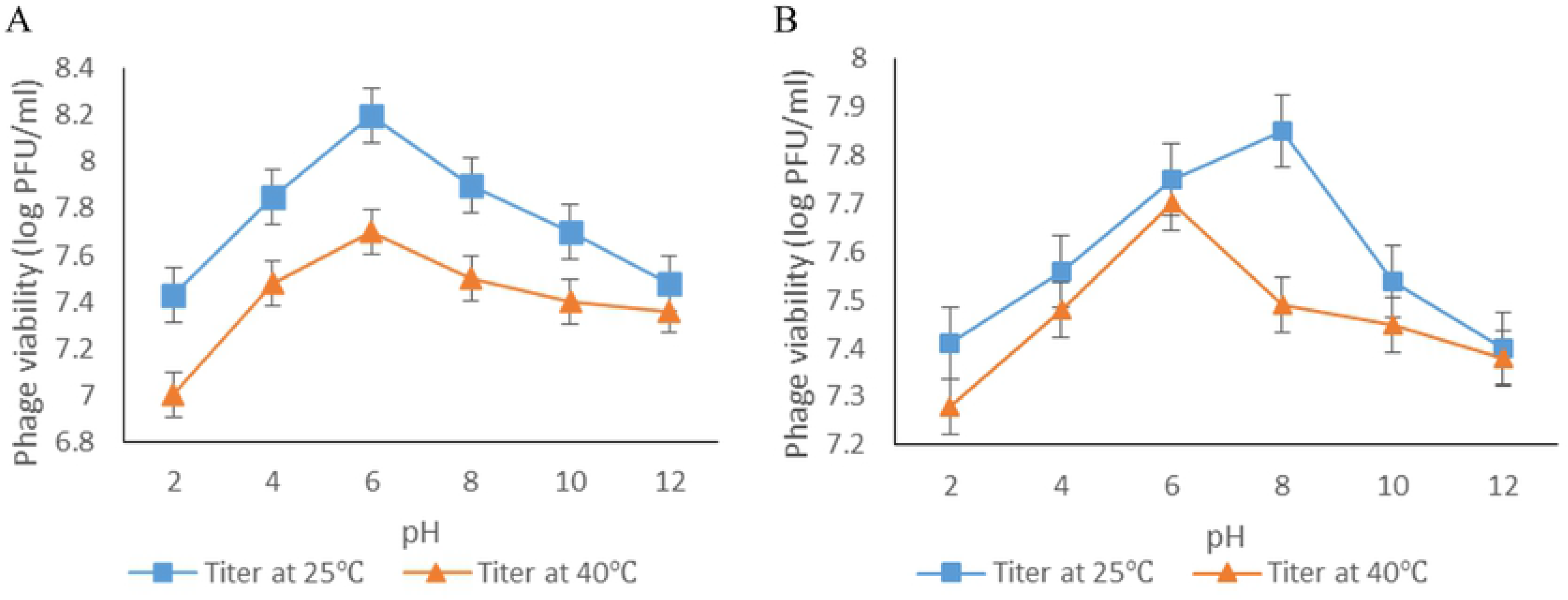
Effect of pH on phage viability at 25°C and 40°C. A) Phage UPEC04. B) Phage UPEC10. Phage viability was determined by obtaining the phage titers at the different pH using the agar overlay method. Values are an average for duplicate tests.

## 4. Discussion

The APEC isolates showed high resistance to commonly used antibiotics in poultry like tetracycline, ampicillin and cotrimoxazole. This high level resistance has been reported in *E. coli* from poultry and other sources [3,28]. This is likely to be as a result of irrational drug use, especially among the poultry farmers and use of antibiotic supplemented feeds. Indeed, in Uganda, Bashahun & Odoch (2015) reported that 96.7% of the poultry farmers frequently used antibiotics for prevention and control of infectious diseases while 33.3% used the antibiotics to promote growth and enhance feed efficiency [29]. Resistance to antibiotics not commonly used in animal production systems, such as chloramphenicol was unexpected but the ease of access without a valid prescription to human drugs over the counter in pharmacies results in their misuse in animals [30]. The latter is likely to be the explanation for the average frequency of resistance that was encountered in case of Chloramphenicol (35.7%). Low frequency of resistance was revealed in case of Gentamicin (10.7%) and Nitrofurantoin (8.9%); while all the 56 (100%) isolates were susceptible to Cefixime. Susceptibility of all the isolates to Cefixime, could be due to the fact that this is a recently introduced antibiotic, quite expensive and not readily available to the farmers. This is in agreement with a study done by Dou *et al* (2015) who found out that there was low resistance towards newly developed drugs [31]. A high rate of multidrug resistance has also been reported elsewhere [10,31,32].The high level of antimicrobial resistance of APEC demonstrated in this study calls for stringent regulations on antibiotic use on poultry farms. Additionally, due to the challenges of developing new antibiotics, the high resistance rates reiterates the need to introduce alternatives to drug use, such as the bio-control agents, like the bacteriophages.

Phylogenetic typing determines the genetic background or ancestry of an organism as well as differentiating between the pathogenic *E. coli* strains (B2 and D) from commensals (A and B1) [33,34]. Overall, phylogenetic analysis of APEC strains in this study revealed that majority belonged to Phylogenetic groups A and D. This is in agreement with several studies done elsewhere [20,28,35,36]. Johnson *et al* (2008) found out that majority of the APEC isolates characterized belonged to A, B1 and D phylogenetic groups [9]. The 11 isolates from group A and the three isolates from group B1 that lacked the virulence genes but were isolated from colibacillosis suspect birds probably harbored other virulence genes that were not tested for during the current study or they were just opportunistic. This is in agreement with Picard *et al* (1999) who found out that some strains of *E. coli* belonging to Phylogenetic groups A and B1 exhibiting commensal characteristics would cause disease [37]. The 14 and five isolates from group A and B1, respectively; that possessed virulence genes could have acquired them by horizontal gene transfer from the pathogenic strains [31,38]. The three isolates from phylogroup D that lacked the tested virulence genes probably caused colibacillosis by possessing other virulence genes not screened for in this study. The above findings agree with other studies that demonstrated diversity of Phylogenetic groups among APEC [39,40].

The selected virulence genes occurred in 69.6% of the *E. coli* isolates with varying frequencies; indicating that they were potentially pathogenic. However, only 50% of the isolates that had four or more genes can be categorized as APEC according to Johnson *et al* [9]. The findings are supported by Kuhnert *et al* who concluded that pathogenicity of a given *E. coli* strain is mainly determined by specific virulence factors which include adhesins, invasins, toxins and capsule [41]. Seventeen isolates (30.4%) did not exhibit a single virulence gene. These isolates could have been commensals that had become opportunistic due to host-dependent factors like other infections, environmental stress, poor nutrition and hygiene [37,39,42]. Alternatively, these isolates could be harboring other virulence genes that were not screened for in the present study [9,32]. Several studies show that it is rare for all the virulence genes to be present in the same isolate [20,31,43]. For instance, Delicato *et al* reported that 27.5% of the colibacillosis-derived isolates did not possess any of the virulence-associated genes investigated [44].

The Episomal outer membrane protease gene (*ompT*) showed the highest prevalence at 58.9%. This gene encodes a protease that cleaves colicin, an inhibitory protein produced by other *E. coli* [45]. The *ompT* gene is located on the ColV plasmid alongside other virulence genes like *iss*, *hlyF* and *iroN* [46]. A relatively high number of isolates harbored the *ompT* gene for protection against colicin produced by other *E. coli*.

The lowest frequency was shown by Salmochelin siderophore receptor gene (*iroN*) at 41.1%. Like the *ompT* gene, *iroN* is located on the ColV plasmid and is one of the genes responsible for iron acquisition [45,46]. Presence of virulence genes distinguishes APEC from commensals and as a result these can be used as molecular markers for detection of colibacillosis in combination with other diagnostic tools [47]. However, there is need to determine whether the various isolates from this study are capable of establishing an infection in order to confirm their pathogenicity.

Out of the 56 APEC isolates, none belonged to the serogroups O1, O2 and O78 which were reported to be the most common [48]. This means that the above serogroups are not common among APEC infecting chicken around Kampala. This can be explained by the fact that distribution of serogroups varies from one region to another and that the APEC serogroups O1, O2 and O78 may not be as common as indicated in other countries like China [20,31]. Indeed, Riaz *et al* reported occurrence of serogroups O1 and O2 but not O78 [49]. Ewers *et al* also demonstrated that colibacillosis can be associated with serogroups other than O1, O2 and O78 [38]. Over 100 APEC serogroups have been reported and most of the previous research was carried out in Europe, Asia and some in Brazil, which are geographically distant from Uganda [20,31,48]. The difference in the prevalent serogroups is not unexpected and infers that vaccines against avian colibacillosis developed elsewhere may not offer protection to chicken in Uganda.

From the findings regarding host range, no single phage was able to lyse all the studied APEC strains. The maximum number that could be lysed was 14 out of 56 (25%). This is because phages are highly specific towards their hosts [50]. This is in agreement with other studies that demonstrated that phages usually have a limited host range [25,51]. Having a relatively broad host range is one of the desirable properties for selection of candidates for phage therapy [52]. The two phages, UPEC04 and UPEC10, which had a combined lytic activity against 14 APEC isolates are better candidates for formulation of cocktails for therapeutic intervention compared to the others. However, there is need to obtain more phages with a wider host range by using either a mix of multiple host strains of the same species for phage isolation or growth on multiple hosts sequentially, that is, one host at a time [52]. Lysis of the eleven multi-drug resistant APEC isolates by the phages demonstrates the potential of phages in controlling infections caused by multi drug resistant bacteria.

The main physical factors affecting phage adsorption and growth include pH and temperature [53]. The different pH and temperature ranges in this study were selected to mimic those that would be encountered during the handling and application of these phages as therapeutic or sanitizing bio-control agents on poultry farms. Both UPEC04 and UPEC10 were stable to heat up to 60°C. At 70°C, the phages were inactivated which is in agreement with Lu *et al* (2003) and Shende *et al* (2017) who reported that phages get inactivated at 70°C and above [51,53].

The effect of pH on phage viability at 25°C and at 40°C represented activity at room temperature and body temperature of chicken, respectively. The two phages were tolerant to a broad range of pH similar to what was observed in previous studies [26]. The tolerance to a broad range of temperature and pH coupled with a wide host range, makes the two phages suitable potential candidates for a cocktail product that can be used as an alternative to antibiotics in the control of APEC infections [27].

## 5. Conclusion

None of the APEC isolates analysed belonged to the most common serotypes O1, O2 and O78 as reported elsewhere. Over 80% of the strains exhibited multi drug resistance against the most commonly used antimicrobials. The *E. coli* isolates belonged to various phylogenetic groups, with the majority belonging to phylogroup A and the minority to phylogroup B2. The selected five virulence genes were present in 69.6% of the APEC isolates at varying frequencies. Of the seven phages that were isolated, two had the highest combined host range of 25% and exhibited lytic activity under a wide range of temperatures and pH, making them potential candidates for a therapeutic cocktail product.

## Recommendations

Future studies can be carried out to determine other virulence genes responsible for the pathogenicity of APEC. There is need to establish the circulating APEC serotypes in Uganda, hence comprehensive screening for other serotypes is necessary. Further investigations are needed to determine the characteristics of the bacteriophages such as, growth rates, latent times, burst sizes, morphology and genome sequences before they can be applied as therapeutic agents in the control of colibacillosis.

## Acknowledgements

Dr. Wilfred Eneku for availing the post mortem tissues from poultry colibacillosis cases for *E. coli* isolation; Dr. Catherine Schouler of the Institut National de la Recherche Agronomique (INRA), France for providing the positive controls used in this study and the poultry farmers where samples for phage isolation were obtained; are greatly acknowledged.

## Authors Contributions

GK – Molecular analyses, bacteriophage evaluation; PK – Bacteria and bacteriophage isolation; RA – Antimicrobial sensitivity testing, bacteriophage evaluation; SA – bacteriophage evaluation; AN – Molecular analyses; JLN – Conceptualized the research, data analysis. All authors read and contributed to the drafting of the manuscript.

## Availability of data and materials

Relevant data generated or analyzed during this study are included in this article and supporting information files.

## Ethics approval and consent to participate

Not applicable.

## Consent for publication

Not applicable.

## Supporting information

**S1 Table. Host Range. Sensitivity patterns of the seven phages on the 14 APEC isolates**

**S1 Appendix. PCR protocol for detecting the virulence genes**

**S2 Appendix. PCR protocol for determining phylogenetic groups**

**S3 Appendix. PCR protocol for detection of APEC serotypes**

